# Grades, motivation, and resilience: the role cognitive and non-cognitive traits play in the undergraduate research student selection process

**DOI:** 10.1101/641670

**Authors:** Celeste Suart, Meagan Heirwegh, Felicia Vulcu

## Abstract

The academic research experience is extremely rewarding but is also froth with many challenges, setbacks, and frustrations. The publish-or-perish mentality of a research-intensive academic environment demands a certain skill set in its trainees. This selection criterion begins with the undergraduate research selection process, but what is the ideal skill set of an incoming student entering the research experience? The current landscape on this topic is bleak, with little examination of how students are chosen for these positions. We therefore conducted an analysis examining student selection methods, non-cognitive traits, emphasis on grades and medical school future ambitions. Our findings suggest that the top five student traits valued by principal investigators are: motivation, resilience, hard work, inquisitiveness and honesty. Surprisingly, emphasis on grades as a screening tool decreased as age of laboratory and frequency of publication increased. Additionally, we identified an inverse correlation between student interest in medical school and research supervisor interest in selecting the student for an undergraduate research experience. Taken together, our study culminates in a defined set of skills beneficial for an incoming student at the beginning of their research experience. We feel our findings will greatly facilitate the overall undergraduate student selection process in any academic environment.

## Introduction

Hands-on research experiences are an effective means of enhancing undergraduate education [1]. They have been shown to develop the student’s transferable skills in communication, teamwork, inquiry, and independent learning [2–4]. This is in addition to the student’s gain in research and laboratory techniques, which typically are mentioned less in literature than transferable skill benefits [4-5]. A well-documented impact of completing an undergraduate research experience (URE) is how it influences a student’s career trajectory in science. Students who complete UREs are more likely to pursue graduate education [2], [4], [6]. This increased interest in graduate school is seen in both students who were considering graduate work before their URE and those who did not [7].

Thus, it seems logical to delve further into the criteria Principal Investigators (PIs) use to select undergraduate students for an undergraduate research experience. Prior work by Weiner identified a few undergraduate metrics PIs use to select high-ranking graduate students [8]. “High-ranking” in this context refers to graduate students that have performed very well as perceived by their PIs. In Weiner’s study, prior research experience was found to be one of the largest undergraduate discriminators [8]. Additionally, a positive reference letter, preferably from the PI who supervised the URE, was another well-defined metric. Put together, this study strongly argues for research experiences to be included in the undergraduate student curriculum, especially when the student is aiming for a graduate education.

This leads us to the Undergraduate Research Experience itself. What denotes a beneficial URE? UREs are often analyzed through apprenticeship model theoretical frameworks such as Vygotsky’s zone of proximal development, Lave and Wenger’s communities of practice or Baxter-Magolda’s epistemological reflection model [9–11]. All three of these models touch on the professional socialization aspect of apprenticeship, providing a framework for how students transition from outsider to a member of research culture [4], [5]. Students who become fully immersed in research culture are more likely to experience positive URE outcomes, such as increased interest in graduate school [7]. What about the URE selection process itself? What do PIs look for in their selection of undergraduate research students? The ample research on the URE socialization process benefits and effects on student career paths does not mirror the sparse research published on the URE selection process itself. Though literature in medical education does address the selection process, there has been no comprehensive analysis on the URE selection process in academia [12–14]. Understanding the URE selection process in academia will provide beneficial insight for both undergraduate student candidates and PIs.

We, therefore, analyzed how students are selected for UREs within the Department of Biochemistry and Biomedical Sciences at McMaster University. This was accomplished using in-person semi-structured interviews with PIs. We had three main objectives; ascertain which skills allow students to successfully interview for URE positions, identify which non-cognitive traits PIs associated with excellent or poor quality undergraduate researchers, and determine the effect of student grade point average (GPA) on their suitability as a URE candidate.

## Materials and Methods

### Selection Criteria

The possible interview candidate pool was defined as all faculty, joint and associate members in the Department of Biochemistry and Biomedical Sciences who had undergraduate researchers (URs) in their laboratory for at least one year (n=48). A recruitment email was sent to the department, with a response rate of 21% (n=10). Interviewees had worked at McMaster University for 1-16 years and were in charge of labs that had published between 4-16 papers in the past two years.

### Interview Methods

Interviews were conducted between February and March 2016 in each PI’s office, inside their respective laboratories, or in a communal office area. The interview process was semi-structured, focusing on the UR selection process, qualities of UR students, and the effect of a student’s GPA on the PI’s decision to select them. This was accomplished with five main questions and three follow-up questions. This protocol is available by request to the authors. If a PI was unsure of the meaning of a particular question, the question was rephrased. If the PI was unsure of the meaning of a word in the question, the word was defined for them or synonyms were given. If an answer given for a question was less than one minute in length, PIs were asked to expand on their answers. Interviews took between 10 and 30 minutes. Audio-recorded interviews were transcribed verbatim using REV.com, an audio transcription service based in California. Transcriptions were then uploaded to MAXQDA 12, a qualitative analysis software, and then searched for themes relating to the research focuses. No personal identifying information was collected at any point during the interview process. All information obtained during the study was de-identified and aggregated during analysis. Identification of individuals is not possible after this process.

### Analysis Methods

The method of data collection and analysis used was based on ethnographic, a tradition rooted in anthropology which is used to describe facets of a culture-sharing group [15]. In an ethnography, theories are developed to explain patterns of cultural behaviour, such as ideas and beliefs. Qualitative analysis software was then used to identify multiple patterns by coding the dataset. Although it is rare for statistical significance to be met, results derived from careful analysis through data coding has the potential to be very powerful [5]. For this analysis, text segments mentioning different ideas were labeled using code names. These codes were not preconceived but were created as ideas arose from the text. During the coding process, the codes and their associated text from the transcript create a dataset for each interview. Groups of codes that center around a theme are grouped together by parent codes. Words used for coding and their definitions are continuously updated and stored in the codebook feature of the analysis software. Information is triangulated by testing one source of data against another and verifying internal consistency as it is being coded [9]. This is done to ensure that all content is valid.

The frequency that a code appears in the text, along with what other codes it appears with are analyzed for patterns and trends. Themes are defined in qualitative analysis by codes that frequently cluster together. This can inform us how and if two ideas are linked together in the minds of the interviewees. Relative frequency can also be used to estimate how important a certain subject is to an interviewee. Since these ideas are not taken from a random sampling, but a pre-selected group, we cannot analyze frequencies for statistical significance [5]. However, this can be examined in future studies on this subject.

### Research Ethics

This study was examined and approved by the Hamilton Integrated Research Ethics Board (Project Number 1053). Written informed consent was obtained from all participants in the study. Participants were informed that they could withdraw from the study at any point up to two weeks following their interview and that no identifying information would be collected.

## Results

The general methods used by laboratories to approach the URE application process can be divided into four stages: the initial student contact, request for further documentation such as student transcripts, a primary interview with the PI and a secondary interview with current laboratory members (S1 Fig). General trends indicate email as the main contact method (S1 Fig, A), followed by the use of transcripts/ curricula vitae (S1 Fig, B), an interview conducted by the PI (S1 Fig, C) and a final decision based on the PI with input from the lab (S1 Fig, D).

Seventy percent of PIs examined transcripts for the types of courses students had taken, in addition to grade level (S2 Fig). When examining transcripts, PIs verified that students had taken courses relevant to their area of research and had a minimal amount of non-degree requirement or “easy pass” courses (Fig 1). Overall, easy and unrelated courses on a transcript were perceived as a weaker candidate application, whereas relevant and laboratory skills courses created a stronger application. Additionally, PIs examined transcripts for trends student GPA including improvement and consistency (S2 Fig).

**Fig 1.**
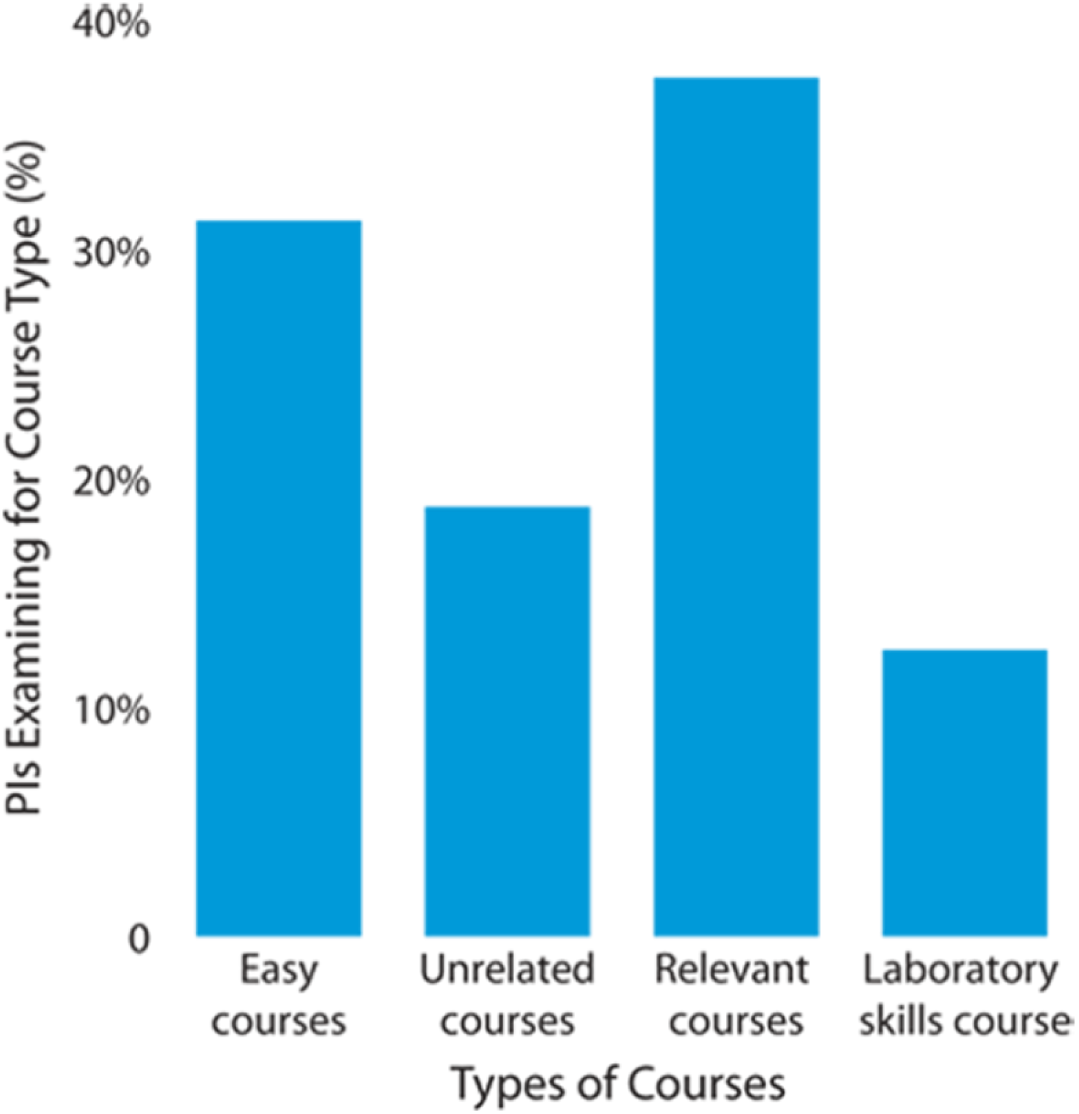
Course types identified on student transcripts by principal investigators. Percentage of PIs interviewed who examine transcripts for specific course types during student selection. Easy courses are colloquially known to have a high class average with minimal student effort. Unrelated courses do not relate to the area of research or the student’s degrees. Relevant courses constitute material directly related to the PI’s area of research.

During the primary interview and document screening, PIs assessed student motivation for applying to a URE. One major question asked in the URE application process was the student candidate’s interest for applying to medical school. Based on this answer alone, ninety-one percent of PIs interviewed expressed a negative opinion of URE candidates considering medical school. Furthermore, the remaining 9% consisted of neutral PI opinions to URE candidates expressing an interest in medical school with no PIs expressing a positive opinion regarding this topic (S3 Fig). This implies a strong inverse correlation between the URE candidate’s motivation for medical school and the PIs interest in selecting said URE student candidate. The purpose of the secondary interview was mainly to assess the candidate’s social interaction, with 90% of PIs saying laboratory social fit was either somewhat or extremely important in candidate selection (S4 Fig).

The concept of “fit”, with respect to a candidate’s fit into a laboratory culture, overarches many traits of which non-cognitive traits play an important role. Non-cognitive traits are a broad range of behaviors and work habits that have a low correlation with conventional tests of cognitive ability [16], [17]. We attempted to analyze non-cognitive traits from two viewpoints: overall value (or ranking) of each trait and a comparison of individual laboratory primary, secondary and tertiary preferences or dislikes with respect to said traits. The most valued non-cognitive traits identified in an excellent undergraduate researcher (UR) were motivation (26.2%), resilience (14.6%), hardworking (13.8%), inquisitiveness (12.3%), honesty (11.5%), good interpersonal skills (9.2%), critical thinking (6.9%) and eloquence (4.6%) (Fig 2A). This order correlated with our analysis of the primary, secondary and tertiary preferences of non-cognitive traits. The primary preference is the trait that is most valued in URs within a particular laboratory. The secondary and tertiary preferences are respectively the second and third most valued trait within URs. The incidence of a trait appearing as either a primary, secondary or tertiary preference across multiple laboratories was then recorded. Motivation was the highest primary preference amongst labs, with resilience being the most common secondary preference and hard work the highest tertiary preference (Fig 2B). The most disliked UR traits were lack of motivation (31.9%), lack of effort (23.6%), lack of resilience (11.1%), poor interpersonal skills (9.7%), lack of critical thinking (8.3%), overconfidence (6.9%), dishonesty (5.6%) and inarticulate (2.8%) (Fig 3A). The most common primary, secondary and tertiary disliked traits amongst laboratories were lack of motivation, lack of effort and poor interpersonal skills respectively (Fig 3B). The meaning of the non-cognitive trait as they pertain to the context of this study are listed below (Tables 1 and 2).

**Table 1.**
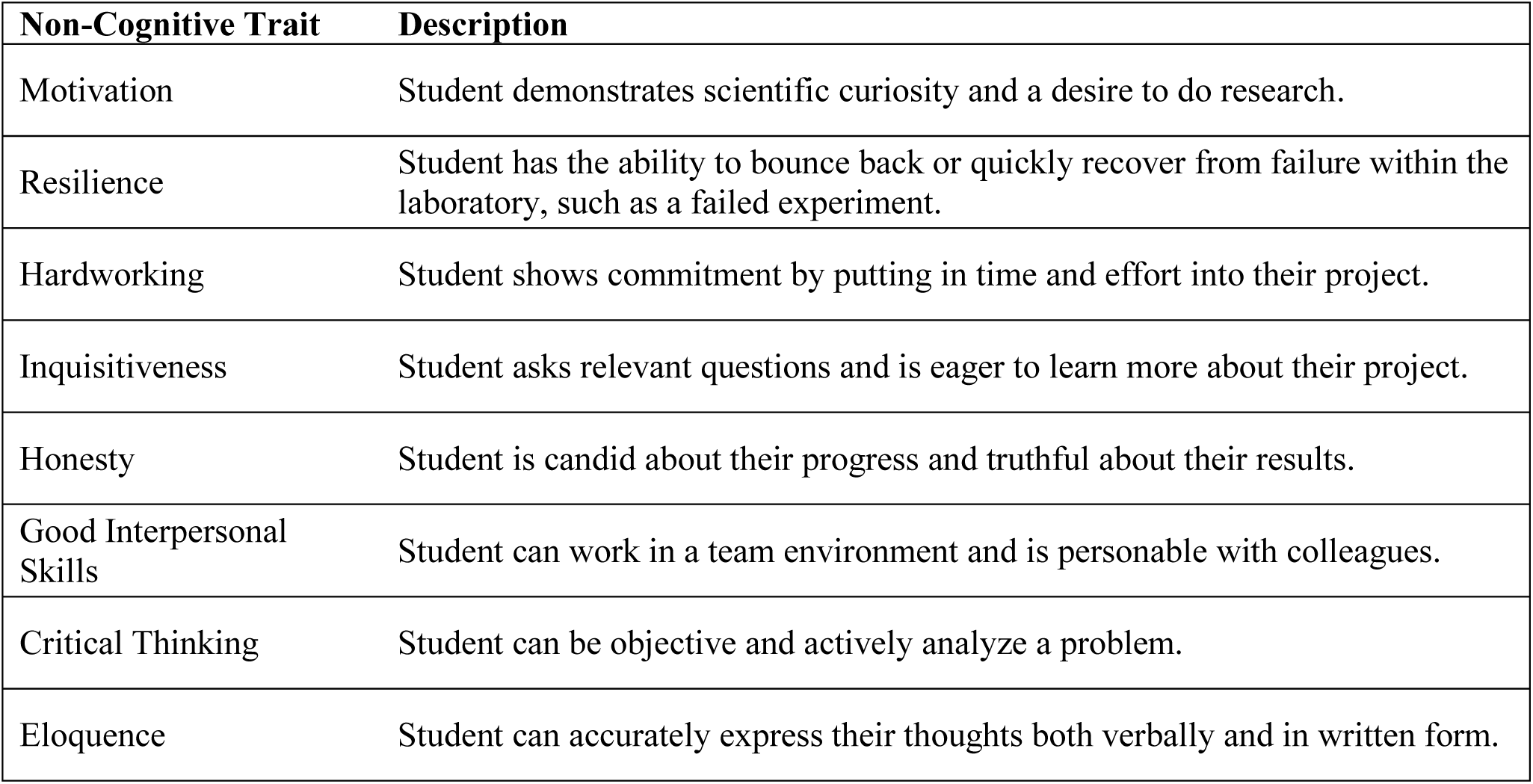
Descriptions of Positive Non-Cognitive Traits.

**Table 2.** Descriptions of Negative Non-Cognitive Traits.

| **Non-Cognitive Trait** | **Description** |
| --- | --- |
| Lack of Motivation | Student is apathetic towards research or is motivated to obtain reference letters instead of a genuine research experience. |
| Lack of Effort | Student puts little or no time and energy into their project. |
| Lack of Resilience | Student has difficulty coping with failure in the laboratory, may not be able to move on to next assignment. |
| Poor interpersonal Skills | Student is difficult to work with on a team and disagreeable with colleagues. |
| Lack of Critical Thinking | Student only examines the surface of problems, performs simple analyses. |
| Overconfidence | Student has an overinflated view of their ability to perform in the laboratory. |
| Dishonesty | Student is guarded about their progress and tends to be misleading about their results. |
| Inarticulate | Student is unable to clearly express their thoughts verbally or in written form. |

**Fig 2.**
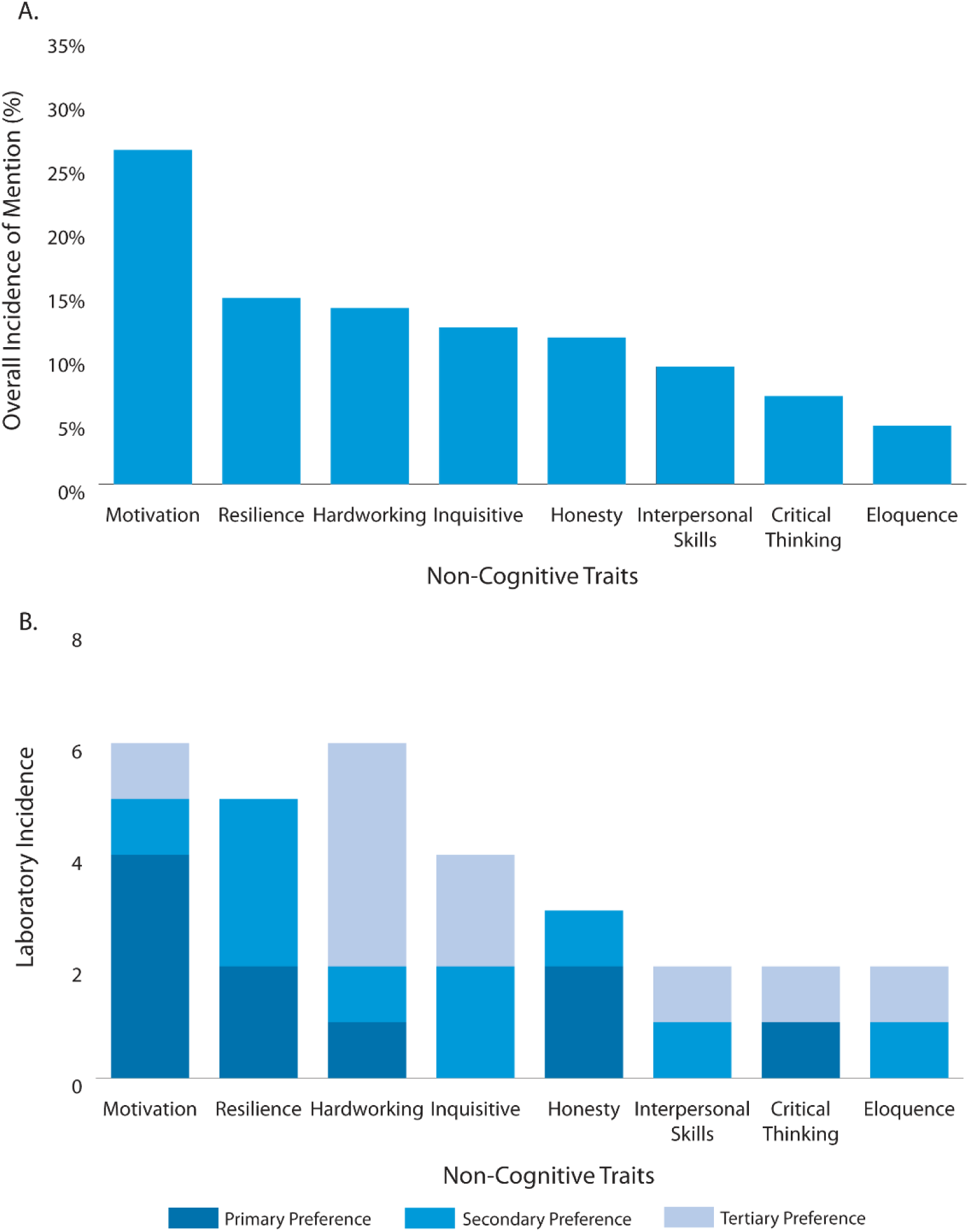
Non-cognitive traits valued in undergraduate student researchers by principal investigators. Traits associated, by PIs, with excellent undergraduate researchers. (A) Overall value ranking of traits based on its prevalence of mention during the entire interview process. (B) Incidence of each skill as the primary, secondary and tertiary preference of each principal investigator interviewed. Quaternary or lower preferences are not included.

**Fig 3.**
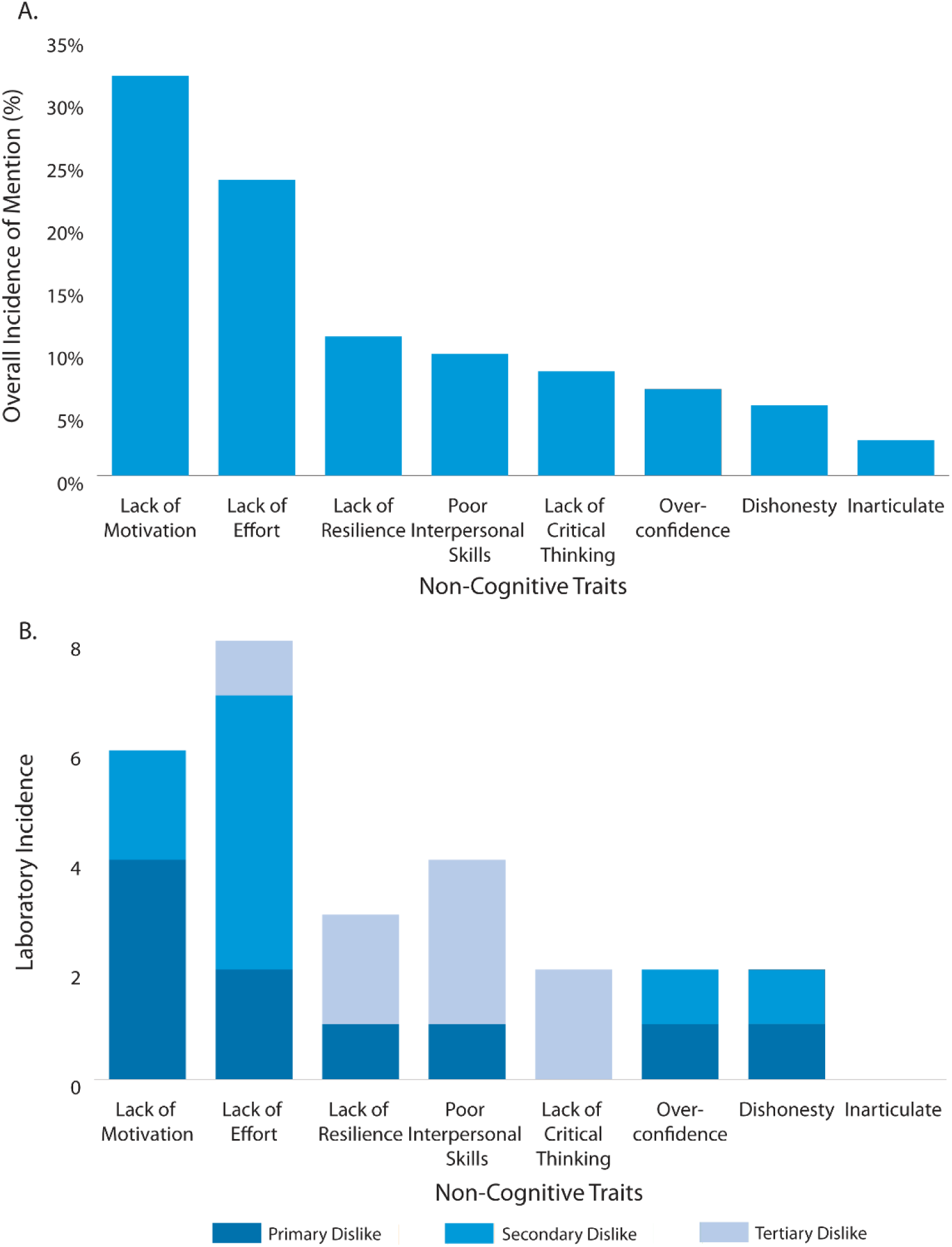
Non-cognitive traits seen as detrimental to undergraduate students by principal investigators. Traits associated by PIs with poor undergraduate researchers. (A) Overall detriment ranking of traits based on the prevalence of mention during the entire interview process. (B) Incidence of each skill as the primary, secondary and tertiary dislike of each principal investigator interviewed. Quaternary or lower preferences are not included.

When asked to describe what traits were typically observed in a lab setting from different caliber URs, PIs mainly described high-quality URs with non-cognitive descriptions and poor quality URs by their lack of technical skills (S5 Fig). This correlated with our previous finding that disliked or negative UR traits were mainly the inability to display a positive non-cognitive trait. It also mirrors how weaker UR applicants were perceived to have more unrelated courses on their transcript than relevant or laboratory skill courses. Without the proper prior training, URs would not have the opportunity to develop technical laboratory skills.

The importance of student grade point average (GPA) was also examined with respect to the age and productivity of each laboratory. Productivity was defined as the number of papers published within the past two years (January 2014 – December 2015). An interesting trend emerged whereby as the number of publications increased, the importance placed on grades by the PI decreased with respect to the importance placed on this selection criterion (Fig 4A, compare upper quartile to lower quartile). A similar trend was seen with laboratory age, whereby grades had less importance as age of laboratory increased (Fig 4B, compare upper quartile to lower quartile). This suggests that PIs with more experience or more resources tend to favour qualitative UR selection criteria while younger PIs tend to use quantitative measures.

**Fig 4.**
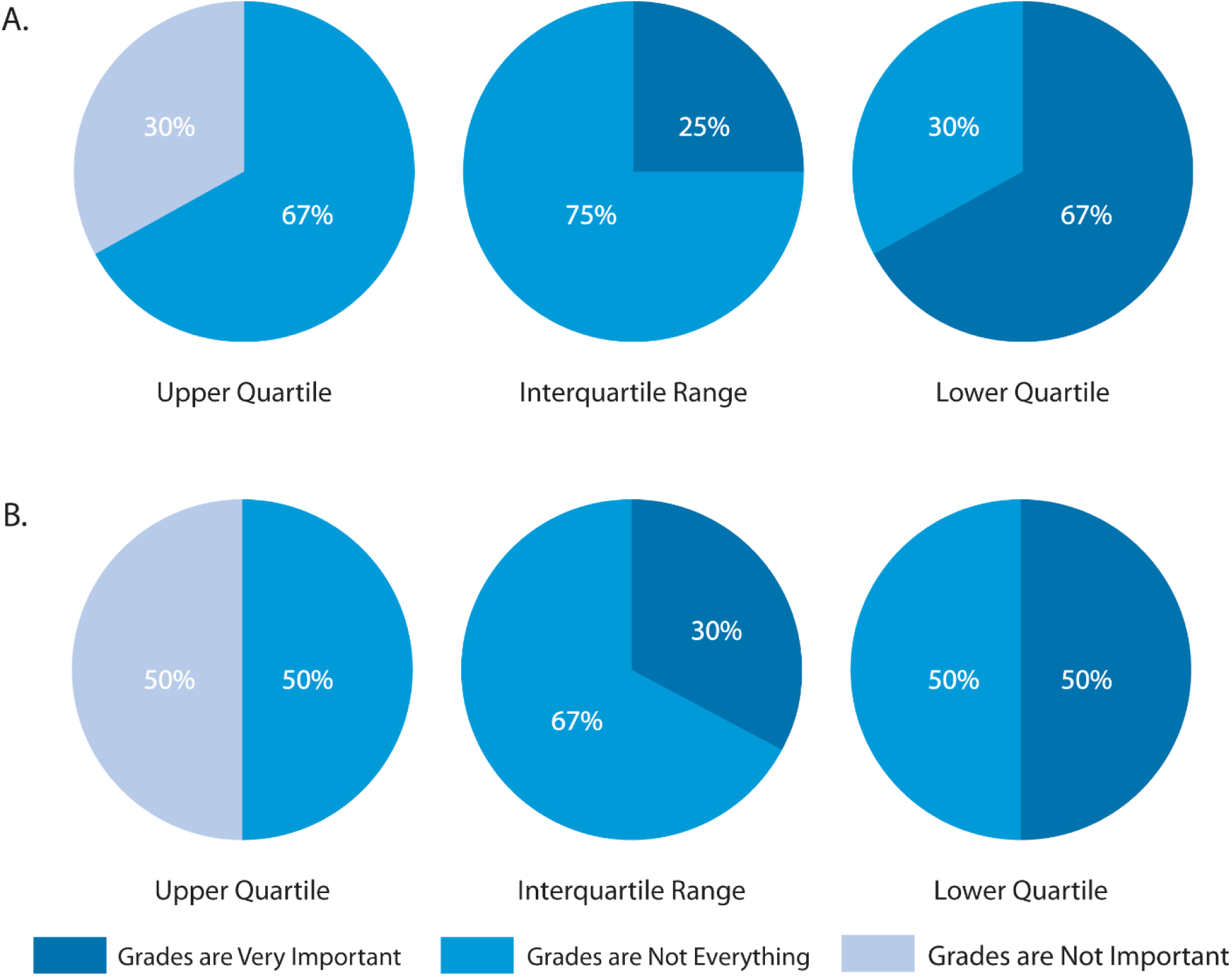
Importance of student grade point average to principal investigators with regard to laboratory productivity and age. All laboratories examined were classified either in the upper quartile, interquartile range or lower quartile in **(A)** lab productivity, represented by the number of articles published between January 2014 - December 2015 and **(B)** the number of years the lab has existed at McMaster University. PIs who viewed grades as “very important” used them as a primary screening tool or placed an emphasis on high grades. PIs who viewed grades as “not everything” placed less emphasis on grades, using them a secondary screening tool. PIs who viewed grades as “not important” did not consider student GPA during the selection process.

Overall, students with the highest GPAs were more often referred to as poor quality (61.9%) than excellent (41.7%) URs (S6 Fig). Students with mid-range GPAs were often referred to as excellent URs (54.2%) while only comprising 4.8% of the poor UR descriptors (S6 Fig). Low GPA students were mainly labeled as poor URs (S6 Fig). Taken together this implies students at both GPA extremes (very high GPA and low GPA) make poorer URs than their mid-range GPA counterparts.

## Discussion

Overall, our study has identified a number of trends which provide a glimpse into the shared departmental beliefs and attitudes that influence student UR selection. Categorizing the URE application process into four stages (Fig. S1) allows us insight into specific features and traits deemed important by the PI throughout this process. The act of looking for specific courses in transcripts suggests PIs prefer students who have given enough forethought to UREs and have planned ahead to take relevant courses (Fig. 1). At first glance this implies student motivation. However, the definition of motivation in this specific study can be further curtailed to specify motivation with respect to research. This was evident from our finding that an indicated interest by the UR candidate in medical school was strongly correlated to a negative bias with respect to said candidate (S3 Fig). These UR student candidates were often seen as seeking reference letters for medical school applications rather than a genuine research experience. This suggests PIs are looking to invest time and effort into students who will continue within academia following the completion of their URE.

Nearly all non-cognitive traits identified on the valued list had their converse trait listed on the detrimental list (Fig 2A and 3A). This observation not only served to validate our method of analysis, but also reiterated a consistent trend in non-cognitive traits coveted by our PIs. For example, motivation, eloquence, and their converses had the same priority ranking on both lists (Fig 2A and 3A). The remaining converse trait pairings fluctuated in priority between the valued and detrimental lists when compared overall and between laboratories. Interpersonal skills were the highest tertiary dislike, yet the converse trait was not in the top three preferences (Fig 2B and 3B). This suggests that the obvious absence of interpersonal skills is more harmful than its presence is helpful to URs. Resilience had a slightly higher priority than lack of resilience, implying its presence is more beneficial than its absence is harmful (Fig 2A and 3A). Inquisitiveness (fourth valued trait) and overconfidence (sixth detrimental trait) did not have a converse trait on the opposite list (Fig 2A and 3A). One possible explanation is their converses, apathy, and insecurity, are easily identified in candidates and screened out due to their level of detriment perceived by PIs.

PI descriptions of different caliber URs emphasizes the importance of candidate non-cognitive traits. The most common descriptions of excellent URs were non-cognitive in nature, such as overall productivity (hard working) or taking pride in a project (motivation) (S5 Fig). Poor quality UR descriptions were mainly skill-based, such as poor troubleshooting or poor execution of lab techniques (S5 Fig). It could be that poor laboratory skills overshadow any valued non-cognitive skill the student may possess. Another possible explanation is that certain non-cognitive traits help in the development of laboratory technique. An example of this could be a high resilience student quickly developing troubleshooting ability since they recover from failure more easily than peers.

Trends in GPA importance, when assessed by lab productivity and age, showed PIs placed less emphasis on student grades as the comparing variable increased (Fig 4). This suggests that emphasizing grades as a screening tool are used most often in smaller or younger labs, perhaps since grades are more quantifiable than non-cognitive traits. Judging students based on quantitative measures is more straightforward and less time consuming than trying to identify candidates through qualitative means. More experience PIs could also have a better idea of what non-cognitive traits they look for in candidates, compared to PIs who have less experience selecting undergraduate candidates.

PI associations of UR quality with previously obtained grades offers some insight into PI assumptions about non-cognitive skill development. There was a prevailing view that students with extremely high averages will have poor lab skills and difficulty coping with challenges faced in the lab (S6 Fig). This implies PIs assume students with high cognitive skills may not have had the opportunity to develop beneficial non-cognitive traits. However, students with mid-range grades, who have presumably struggled more with courses, are seen as more probable to have developed non-cognitive traits such as resilience. This may explain why certain PIs examined trends in student GPA (S2 Fig). A student whose GPA improved over time may have struggled earlier, developed beneficial non-cognitive traits and successfully applied these new skills to improve their average.

The benefits of UREs for students are known both colloquially and within undergraduate education research. However, the selection process for UREs has been largely overlooked as a field of study. As a result, students can often be distressed during URE selections due to lack of prevalent information. This is also detrimental to PIs, who must use trial and error to adjust their UR selection method, which is a tedious and time intensive process. This work begins to fill this gap of candidate selection in URE literature.

These findings can aid both undergraduate students and PIs by clarifying the URE selection process. Although these results are specifically related to the departmental culture examined, this same method could be applied to other departments and universities to identify aspects of research culture specific to those cultures. We envision our study to be an important step towards understanding the URE selection process.

## Acknowledgements

Interviews, data analysis, and manuscript preparation were conducted by Celeste Suart. The authors thank the faculty members in the Department of Biochemistry and Biomedical Sciences (McMaster University, Hamilton, Canada) who volunteered their time to participate in this study.

