## Supplemental data figures for "Grades, motivation, and resilience: the role cognitive and non-cognitive traits play in the undergraduate research student selection process"

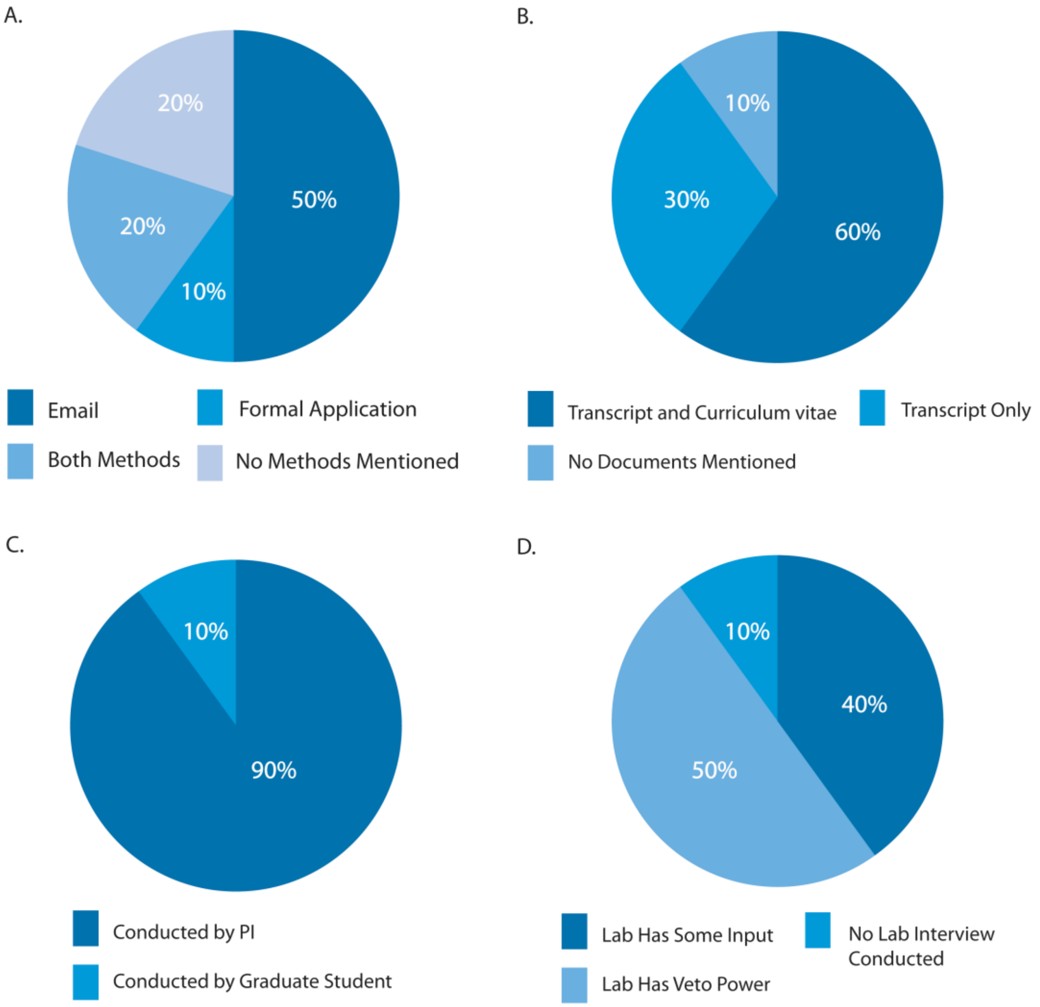


**S1 Fig. Differences in laboratory approaches to the URE candidate selection process.**

Percentage of laboratories using each method shown for each stage. (A) Contact method between URE candidates and principle investigator (PI). (B) Student documents requested by PIs. (C) Identity of the primary interviewer. (D) Incidence and importance of group laboratory interviews.

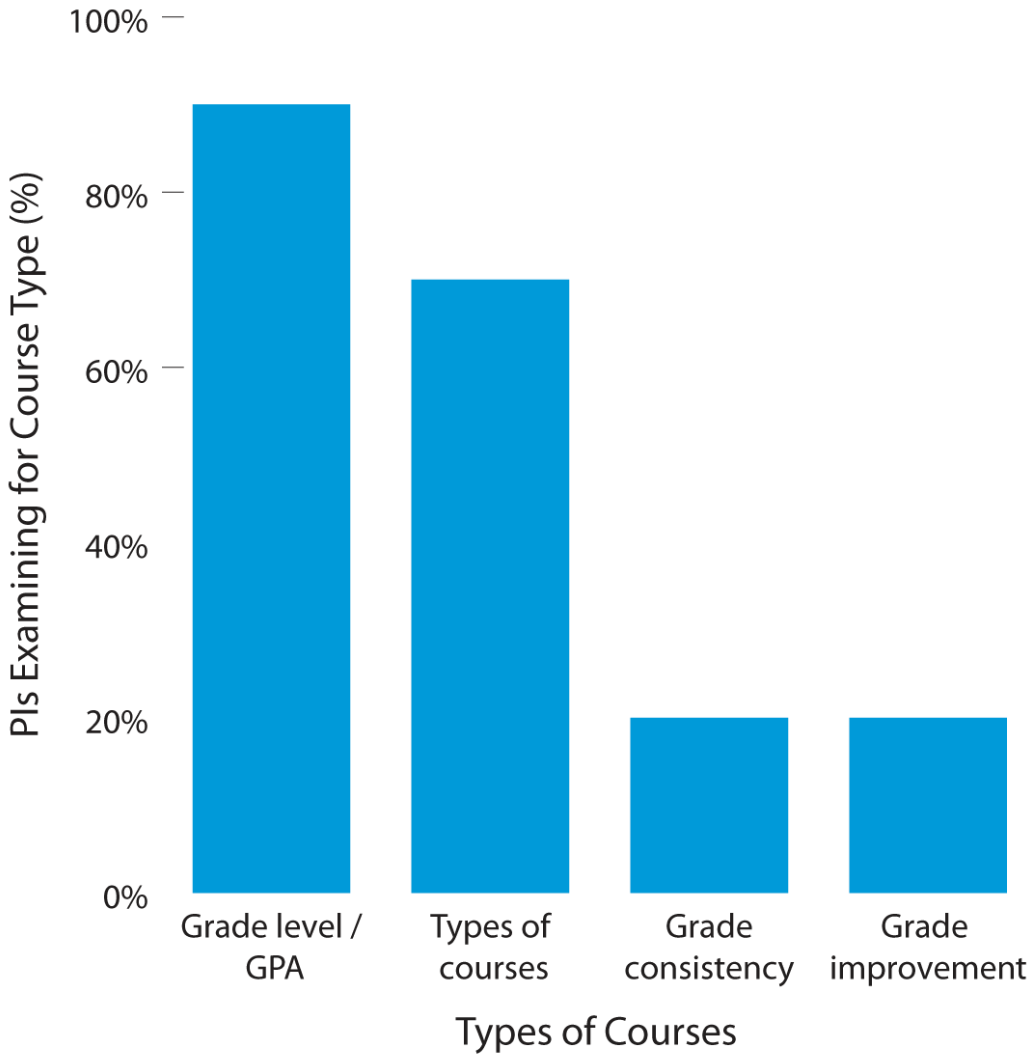


**S2 Fig. Student transcript information examined by principal investigators during the URE candidate selection process.** Percentage of PIs examining transcripts for overall grade level, type of courses, student grade consistency and grade improvement over time.

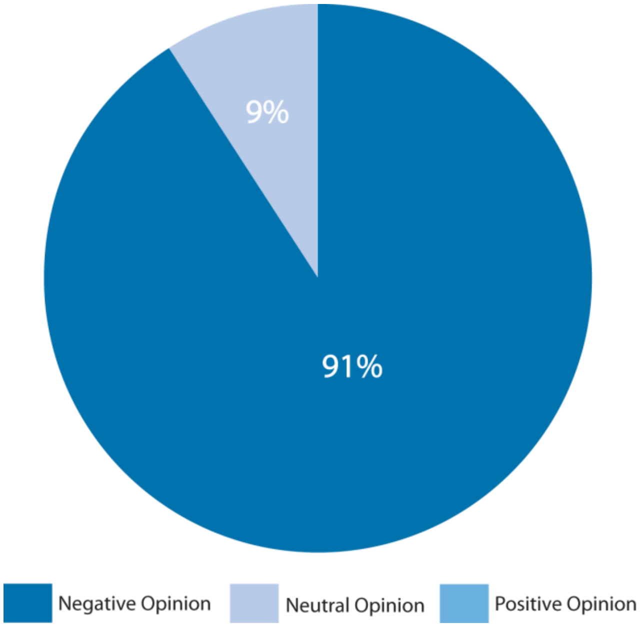


**S3 Fig. Principal investigator opinions of URE candidates considering applying to medical school.** Percentage of negative and neutral opinions expressed over the course of the study. No positive opinions were documented.

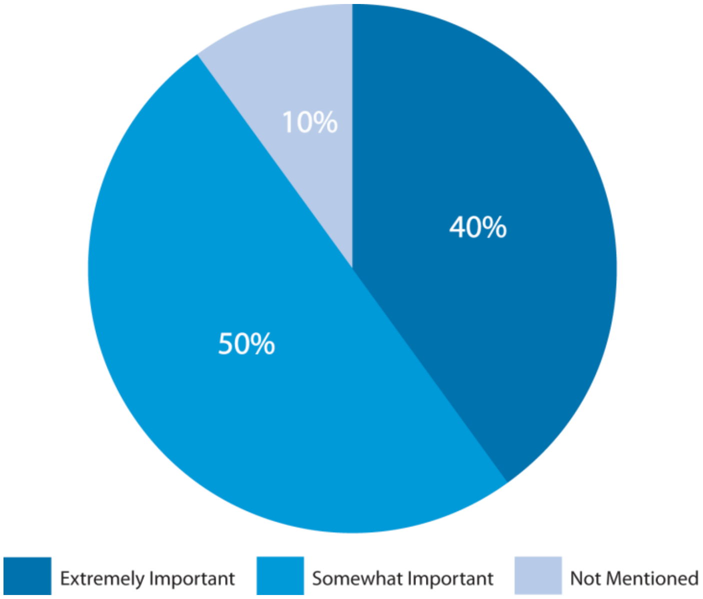


**S4 Fig. Importance of laboratory social fit during URE candidate selection.** Percentage of PI views of how important social fit is during candidate selection. Extremely important indicates that PIs interviewed would not choose a student who was a poor social fit. Somewhat important indicates that poor social fit would make them more hesitant but would consider the student.

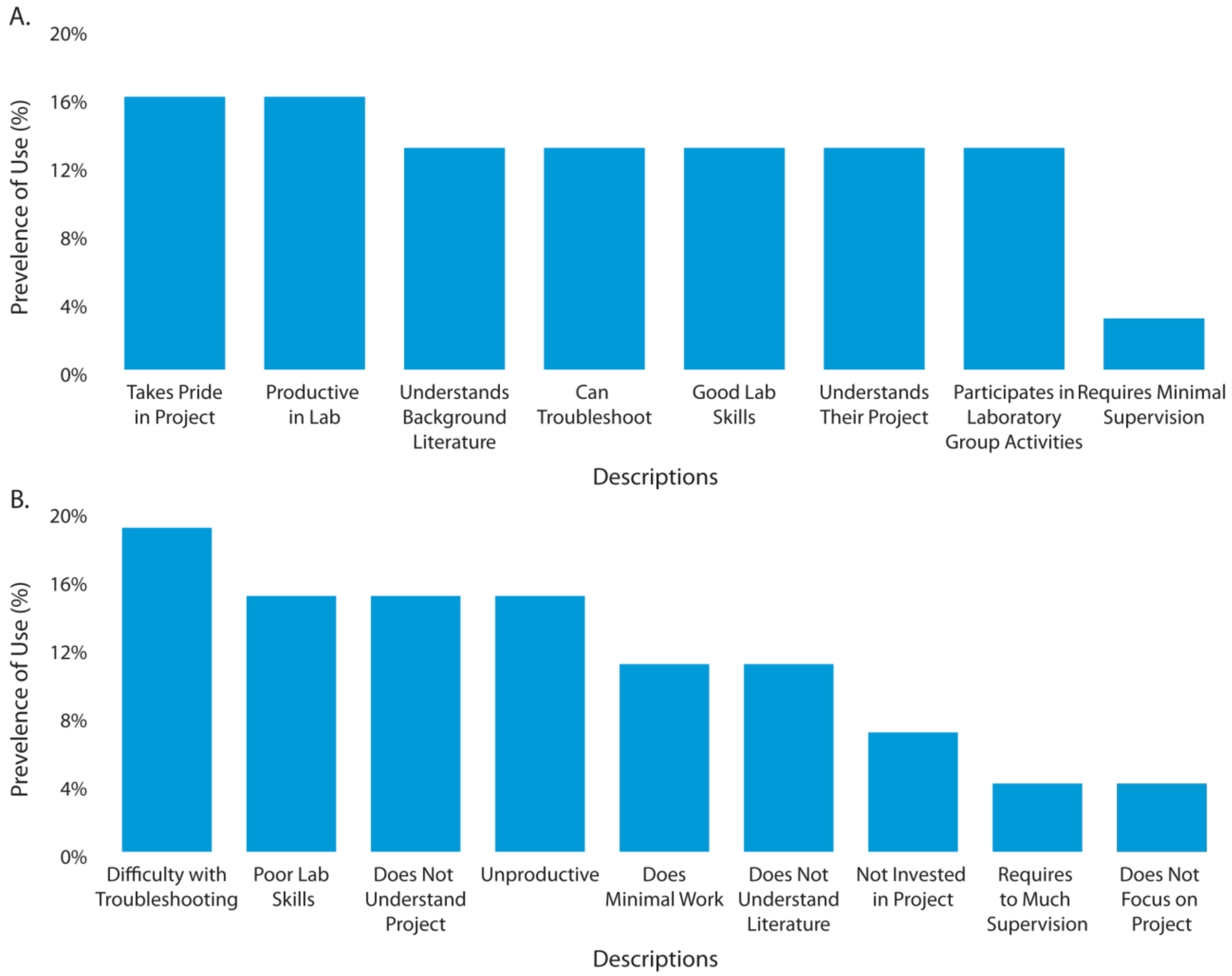


**S5 Fig. PI descriptions of different calibers of undergraduate researchers.** Proportion of descriptions mentioned of (A) excellent and (B) poor undergraduate student researchers based on abilities and behaviours shown in the lab.

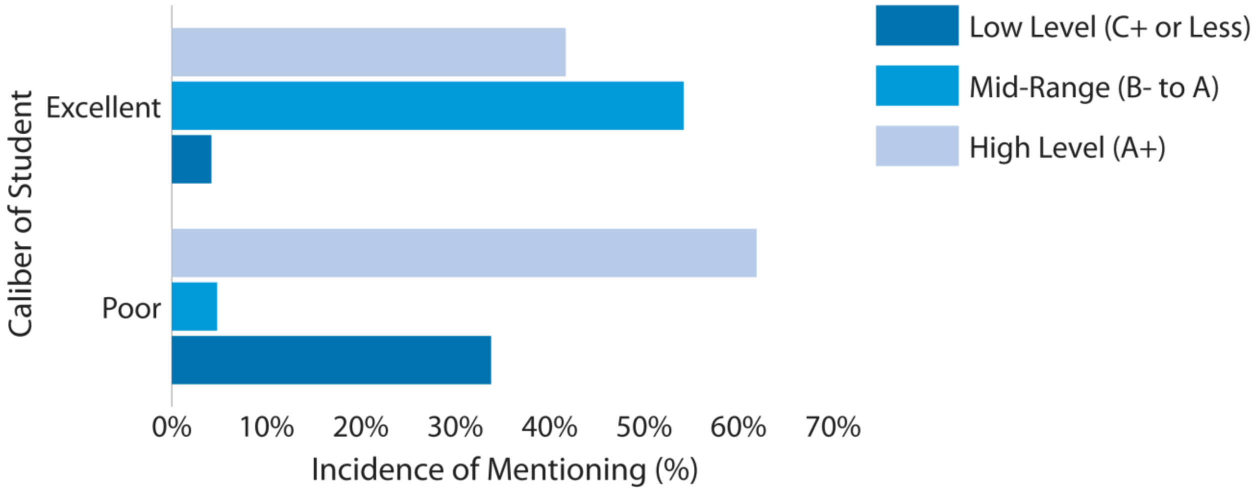


**S6 Fig. PI description of undergraduate student researchers in relation to student grade point average.** All descriptions included a reference to student GPA and judgement of their quality as undergraduate researchers.
